# Development of a Humanized Mouse Model for Studying adult Spinal Cord myelination, remyelination and Drug Efficacy

**DOI:** 10.64898/2026.03.11.711008

**Authors:** Nadjet Gacem, Sabah Mozafari, Jeremy Chazot, Marion Levy, Ana Belén Martínez-Padilla, Radmila Panic, Farina Windener, Gianvito Martino, Tanja Kuhlmann, Brahim Nait-Oumesmar, Anne Baron-Van Evercooren, Beatriz García-Díaz

**Affiliations:** Sorbonne Université, Paris Brain Institute – Institut du Cerveau (ICM), Inserm, CNRS, Hôpital Pitié-Salpêtrière, F-75013 Paris, France; Department of Clinical Neurosciences and National Institute for Health Research (NIHR) Biomedical Research Centre, University of Cambridge, UK; IBIMA Bionand Platform Biomedical Research Institute, (IBIMA)-Plataforma Bionand, Malaga, Spain; Institute of Neuropathology, University Hospital Münster, Germany; IRCCS San Raffaele Hospital, Neuroimmunology Unit, Division of Neuroscience, Milan, Italy; Department of Human Physiology, Human Histology, Anatomical Pathology and Physical and Sports Education, Faculty of Medicine, University of Malaga, Malaga, Spain

## Abstract

Oligodendrocytes are essential for central nervous system (CNS) function through their roles in myelination and neuronal support. Remyelination, the regeneration of myelin after damage, often fails in human demyelinating diseases, leading to progressive neurological dysfunction. While the biology of rodent oligodendroglia has been investigated in depth, species-specific differences have hindered the study of human oligodendrocyte progenitor cells maturation using current animal models. To address this gap, we developed a humanized oligodendroglia chimeric mouse model by transplanting human iPSC-derived O4+ oligodendroglial progenitors (hiOLs) into the developing spinal cord of myelin deficient-immunosuppressed mice followed by demyelination of the adult humanized spinal cord. This paradigm allows in vivo investigation of hiOL fate during their maturation and ageing, and the response of their settled progeny to demyelinating injury.

Following transplantation during postnatal development, hiOLs proliferated, migrated, and differentiated into mature oligodendrocytes, initiating axonal myelination. A population of undifferentiated, proliferative adult human oligodendrocyte progenitors presisted, indicating a reservoir of these cells capable of responding to injury. Spinal cord focal demyelination via lysolecithin injection in the adult humanized spinal cord led to transient loss of human-derived mature oligodendrocytes and myelin, followed by myelin recovery together with endogenous cells.

Treatment with the histamine receptor H3 antagonist, bavisant, a recently identified promyelinating compound, significantly enhanced hiOL maturation and myelin production, indicating promotion of their differentiation when administered post-demyelination. Electron microscopy showed an increased number of human-derived remyelinated axons together with decreased g-ratios in bavisant treated animals.

This humanized spinal cord model of myelination/demyelination-remyelination faithfully recapitulates key aspects of human oligodendrocyte biology and CNS repair, providing a powerful tool to study human-specific myelination and remyelination mechanisms and screen for emerging remyelinating therapies. Its application in disease modeling and preclinical testing holds promise for advancing personalized treatments in demyelinating and neurodegenerative disorders.

## Introduction

Oligodendrocytes play a critical role in the central nervous system (CNS) by producing the myelin sheath, a multilayered lipid-rich membrane that wraps around axons (Stadelmann et al., 2019). The myelination process is essential for the rapid and efficient conduction of electrical impulses along nerve fibers, enabling proper communication between neurons and determining the speed of the nerve impulse (Hughes and Appel, 2016). Beyond insulation, oligodendrocytes also provide metabolic and trophic support to neurons, contributing to their long-term survival (Philips and Rothstein, 2017). Oligodendrocyte damage or dysfunction leads to hypo- or demyelination and severe neurological deficits (Han et al., 2022; Horiuchi et al., 2024; Lopez-Muguruza and Matute, 2023), highlighting their indispensable role in maintaining CNS integrity and function (Mot et al., 2018).

Oligodendrocyte dysfunction is increasingly recognized as a key contributor to the pathogenesis of various neurological disorders. In demyelinating diseases such as multiple sclerosis, impaired oligodendrocyte function leads to myelin loss and axonal degeneration, causing motor and sensory deficits (Falcao et al., 2018). In neurodegenerative conditions such as amyotrophic lateral sclerosis (Kang et al., 2013), Alzheimer’s(Depp et al., 2023) and Parkinson’s diseases (Barba-Reyes et al., 2025), early oligodendrocyte stress and myelin abnormalities exacerbate neuronal vulnerability. Additionally, in leukodystrophies and several psychiatric disorders, genetic and epigenetic disruptions of oligodendrocyte development and function result in white matter defects and neuronal damage (Chen et al., 2018; Fodder et al., 2023; Rey et al., 2024; Schuster et al., 2022). Moreover, oligodendrocyte dysfunction in new myelin formation has been associated with behavioral and cognitive impairments (Falkai et al., 2023; Filley, 2021). These findings highlight the central role of oligodendrocyte health in maintaining CNS integrity across a broad spectrum of pathologies.

Oligodendrocyte progenitors (OPCs) differentiation into oligodendrocytes and myelin formation are regulated by several intrinsic and extrinsic factors (Adams et al., 2021; Boulanger and Messier, 2014). Oligodendrocyte maturation is influenced by the external in vivo environment, including diverse signaling molecules and neuronal activity (Baydyuk et al., 2020; Hill et al., 2025). Intrinsic factors, such as species, tissue and age heterogeneity, control oligodendrocyte internal machinery and are determined by their genetic or epigenetic information (Dimovasili et al., 2023; Howng et al., 2010; Xu et al., 2020). In this respect, several studies highlighted differences in human oligodendroglial cell biology over rodents including their developmental generation period which is 45 times longer than for rodents, the transcriptional regulation and response to growth factors for differentiation, some of which are reflected in vitro and ex vivo in transplantation paradigms (for review (Chanoumidou et al., 2020)). Therefore, to increase the translational relevance of research on oligodendrocyte maturation and myelination, studies should be conducted not only on rodents but also on human-derived oligodendrocytes whenever possible. Intrinsic cues of human oligodendrocyte differentiation can be partially studied in vitro, i.e. in cells cultured with nanofibers (Luo et al., 2022) and ex vivo in organotypic cultures (Tsarouchas et al., 2025). However, (re)myelination is highly influenced by a variety of dynamic extrinsic cues occurring in vivo. Therefore, the development of in vivo humanized models stands out as an essential and urgent need to properly dissect the internal mechanistic of human oligodendrocyte maturation and myelination process occurring in vivo where extrinsic factors have an impact. Investigating specific genetic characteristics within an in vivo environment is crucial for achieving a more precise understanding of the mechanisms of myelin formation. In this regard, devising a humanized model based on human induced pluripotent stem cell (hiPSC)-derived oligodendroglial cells presents substantial translational promise. It may also facilitate the development of personalized therapeutic strategies specifically tailored for individual patients suffering from dys- and demyelinating diseases.

Beyond neurodevelopment, CNS injury or neuroinflammation can damage oligodendrocytes. Under these circumstances, adult OPCs and mature oligodendrocytes are activated to initiate remyelination and proceed to recovery of the altered CNS function (Jeffries et al., 2021; Patel et al., 2010). Although both myelination and remyelination involve myelin formation, remyelination is a reparative process that occurs after myelin damage. While it involves the recruitment and differentiation of adult OPCs and reactivation of surviving oligodendrocytes, remyelination is often less efficient than developmental myelination and results in thinner, structurally different myelin sheaths (Orthmann-Murphy et al., 2020; Sim et al., 2002). Furthermore, mechanisms of developmental myelination and remyelination might involve distinctive challenges (Moyon et al., 2017), as the latter occurs within distinct cellular contexts, ageing, changes in extra-cellular matrix and chronic inflammation (Hill et al., 2025).

In pathological conditions, distinguishing between these intrinsic and extrinsic factors is critical to identify new therapeutic avenues to preserve or mantain oligodendrocyte health and CNS function. Previous studies based on transplantation of human neural precursors (NPC) or OPCs (whether derived fetally, or from induced pluripotent stem cells) in the developing or in the demyelinated rodent CNS highlighted the slow process of differentiation of these human-derived cells and their capacity to myelinate host CNS axons (Ehrlich et al., 2017; Garcia-Leon et al., 2020; Mozafari et al., 2020; Windrem et al., 2020). However, none of these studies analyzed the capacity of hiPSCs-derived progenitors to participate to myelin repair after in vivo aging as adult OPC or myelinating oligodendrocytes. Here, we designed a humanized mouse model that enables to study the fate of human induced O4+ oligodendroglial cells (hiOL) grafted into the developing spinal cord of MBP- and immune-deficient shiverer/Rag2^-/-^ mice. This paradigm allows the grafted cells to mature as adult OPCs and oligodendrocytes during developmental myelination, and to respond as already settled mature oligodendrocytes and OPCs to a demyelination insult. Moreover, we investigate the power of a newly described pro-myelinating agent (Gacem et al., 2026) to enhance the exogenous repair process in this human oligodendroglia chimeric mouse model. This paradigm provides compelling evidence that it could serve as a powerful tool for translational research in neurodegenerative diseases. First, it provides an in vivo system to study the dynamics of human OPCs and oligodendrocytes of a given specific genetic background (e.g. hiPSCs-derived OPC from any patient) to face myelination and to identify any associated abnormalities. Second, it allows analyzing these settled OPC and oligodendrocyte responses to remyelinate after a demyelinating insult. Lasty, this humanized myelination/demyelination-remyelinating paradigm enables to evaluate the therapeutic capacity of any putative pro-myelinating drug on adult human oligodendroglial cells challenged with demyelination.

## Methods

### Animals

The established Shi/Shi:Rag2^−/−^ dysmyelinating-immunodeficient mouse line was chosen as host in order to (i) prevent immune rejection of transplanted human cells and enabling the detection of donor-derived wild-type myelin, and (ii) study the intrinsic behavior of OPCs/Oligodendrocytes derived from hiPSCs in an environment devoid of B and T lymphocytes. All mice were maintained under standard housing conditions, with a 12-hour light/12-hour dark cycle and free access to dry food and water, within the ICM animal facility. All experimental procedures were conducted in accordance with European Community regulations and approved by the INSERM ethical committee (authorization 75-348; April 20, 2005) and local Darwin ethical committee.

### Human cells

Fibroblasts were obtained with informed consent from several individuals. Reprogramming into induced pluripotent stem cells (iPSCs) was performed using a replication-incompetent Sendai virus kit (Invitrogen) following the manufacturer’s instructions. The study received approval from the local ethics committees of Münster and Milan (AZ 2018-040-f-S and Banca INSpe, respectively).

Human iPSCs were differentiated into NPCs using small molecule protocols as previously described (Ehrlich et al., 2015; Reinhardt et al., 2013). NPCs were subsequently directed into oligodendroglial lineage cells expressing O4 by transduction with a polycistronic lentiviral vector encoding the human transcription factors Sox10, Olig2, and Nkx6.2 (SON), followed by an internal ribosome entry site-puromycin N-acetyltransferase (IRES-pac) cassette to enable 16-hour puromycin selection (Ehrlich et al., 2017).

Briefly, NPCs were seeded at a density of 1.5 × 10^5^ cells per well in 12-well plates, allowed to adhere overnight, and transduced with SON lentiviral particles in the presence of protamine sulfate (5 μg/ml) in fresh NPC medium. After thorough washing, the viral medium was replaced with glial induction medium (GIM). Four days later, GIM was substituted with differentiation medium (DM). After 12 days of differentiation, cells were dissociated with Accutase for 10 minutes at 37°C, washed with phosphate-buffered saline (PBS), and resuspended in PBS containing 0.5% bovine serum albumin (BSA). Singularized cells were filtered through a 70-μm cell strainer (BD Falcon), incubated with a mouse IgM anti-O4-APC antibody (Miltenyi Biotech) according to the manufacturer’s protocol, washed, and resuspended in PBS/0.5% BSA at a concentration of 5 × 10^6^ cells/ml. O4-positive cells were sorted using a FACS Aria cell sorter (BD Biosciences), cryopreserved, and stored in liquid nitrogen. Media formulations are detailed in reference (Ehrlich et al., 2017). Human O4+ hiOLs were characterized and their functionality as myelin forming cells was previously validated by engraftment into the neonatal demyelinated spinal cord of immune- and MBP-deficient shiverer/rag2^-/-^ mice (line C1) (Ehrlich et al., 2017; Mozafari et al., 2020).

### Cell transplantation

To investigate the fate of hiOLs during development, cells were transplanted in the spinal cord of three-week-old Shi/Shi:Rag2^−/−^ dysmyelinating-immunodeficient mice. To this aim, mice were anesthetized via isofluorane inhalation (3-4% for induction and 1-3% for maintenance) and local anesthesia was induced by lidocaine prior to the procedure, at the incision site. Each animal received a single, slow injection (1 μl over 2 minutes) of hiOLs (1 μl containing 10^5^ cells/μl) at the thoracic level of the 13th thoracic vertebral level, using a stereotaxic apparatus equipped with a micromanipulator and a Hamilton syringe. Mice were sacrificed at 8, 12, 17 weeks post-grafting (wpg) for subsequent immunohistological analyses.

To investigate the fate of the aged O4+ hiOLs progeny in repair conditions, eleven-week old Shi/Shi:Rag2^−/−^ dysmyelinating-immunodeficient mice engrafted with hiOLs for 8 weeks post graft (wpg) were anesthetized via isofluorane inhalation (3-4% for induction and 1-3% for maintenance) and local anesthesia was induced by lidocaine prior to the procedure, at the incision site. Focal demyelination was induced by stereotaxic injection of 1 μl of 1% lysolecithin (Sigma-Aldrich) (LPC, 1% in 0.9% NaCl) into the spinal cord dorsal funiculus at thoracic level T13 (Blanchard et al., 2013; Buchet et al., 2011). Humanized mice were euthanized at 1 and 8 weeks post-lesion (wpl) for subsequent immunohistological analyses.

### Pharmacological treatment

Bavisant (30mg/kg) in vehicle (0.5% methocel) or vehicle was administered 5 days post LPC injection by daily oral gavage until sacrifice at 8 wpl for immunohistological studies and electron microscopy.

### Immunohistological analyses

Animals were sacrificed (n = 3-6 per condition and time point), via transcardial perfusion with 1× PBS followed by 4% paraformaldehyde in 1x PBS. Following dissection, spinal cords were post-fixed for 1 hour in the same fixative, then incubated overnight in 20% sucrose in 1× PBS.

Spinal cord thoracic segments (∼12 mm, centered on the lesion site) were transversely sectioned into 3–5 pieces (∼3 mm each), serially ordered, and placed in small plastic containers. Samples were embedded in cryomatrix (Thermo Scientific), frozen in cold isopentane at –60°C, and stored at –20°C until further processing. Transverse cryosections (12 μm) were prepared using a Leica cryostat, collected in three series of 10 slides each (30–50 sections per slide), and subsequently processed for immunohistochemistry.

Transplanted cells were identified using human specific anti-nuclei or anti-cytoplasm (1:100; STEM101; STEM 121, Takara, Y40400, IgG1) antibody. The phenotypic characterization of STEM-positive cells was performed with the following primary antibodies: anti-OLIG2 (AB9610, Millipore), anti-CC1 (OP80, Millipore), anti-Ki67 (556003, BD Biosciences), anti-MBP (AB980, Millipore), and myelin anti-MOG (mouse IgG1 hybridoma, clone C18C5; provided by C. Linnington, University of Glasgow, UK). For MBP and MOG detection, tissue sections were pretreated with ethanol for 10 minutes at room temperature. Primary antibodies were revealed using appropriate species-specific secondary antibodies from Alexa Fluor-conjugated antibodies (Invitrogen), combined with Hoechst dye (1 mg/ml) for nuclear counterstaining. Imaging and cell visualization were conducted using a Carl Zeiss microscope equipped with the ApoTome.2 system for optical sectioning and tissue scanning and processed with National Institutes of Health ImageJ software (Schneider et al., 2012).

### Electron microscopy analysis

Mice (n = 3-4 per condition) were transcardially perfused with 20 mL of a fixative solution composed of 4% paraformaldehyde and 5% glutaraldehyde in phosphate buffer (Electron Microscopy Sciences). Following perfusion, spinal cords were carefully dissected, rinsed in phosphate buffer, and the thoracic level cut into segments approximately 700 µm thick. These tissue samples were then post-fixed in 2% osmium tetroxide (Electron Microscopy Sciences, EM-19150) for 1 hour at room temperature.

Samples were dehydrated in graded alcohol before being embedded in epoxy resin. Semi-thin transverse sections (∼500 nm) were stained with toluidine blue to facilitate the identification of thoracic tissue blocks containing the lesion. From the selected blocks, ultrathin sections (∼70 nm thick) were prepared using an EM UC7 ultramicrotome. These sections were subsequently contrasted with uranyl acetate and analyzed using a HITACHI 120 kV HT-7700 electron microscope, allowing high-resolution imaging of ultrastructural features.

The proportion of remyelinated axons in the dorsal funiculus was quantified in bavisant-treated versus vehicle-treated mice, specifically focusing on axons with a diameter ≥ 1 μm (axon numbers, n=1458 in the vehicle group and n= 2364 in the bavisant group). The presence of compact myelin within spinal cord lesions of shiverer mice was used as a criterion to identify myelin derived from the grafted hiOLs. To evaluate the effect of treatment on graft-derived myelination, myelin thickness was quantified by calculating the g-ratio (axon diameter / fiber (axon + myelin) diameter) in the bavisant-treated and vehicle-treated groups using ImageJ software.

### Statistical Analysis

Data are expressed as mean ± Standard Error of the Mean (SEM). Normal distribution was assessed according to the Shapiro-Wilk and Kolmogorov-Smirnov tests. Statistical significance for comparisons between two groups was assessed using the unpaired t test. For comparisons involving multiple groups, one-way analysis of variance (ANOVA) was employed, followed by Tukey’s multiple comparisons tests. All statistical analyses were performed using GraphPad Prism versions 8.0.1 (GraphPad Software Inc., USA). The specific statistical tests applied to each experiment are indicated in the respective figure legends.

## Results

### hiOLs engrafted into the developing spinal cord timely colonize, differentiate into oligodendrocytes and myelinate the spinal white matter and grey matter

To develop a humanized oligodendroglia chimeric mouse model, O4^+^ hiOLs were grafted into the dorsal funiculus of 3-week-old MBP- and immune-deficient shiverer/Rag2^-/-^ mice. This model was selected to provide an immunologically permissive environment for human cell engraftment as well as the capacity to assess graft-derived myelination in the presence of uncompact and thinly-myelinated host axons. Immunohistochemical evaluations were performed at 8-, 12-, and 17- weeks post-graft (wpg) to assess the capacity of grafted human cells to proliferate, colonize, and mature into human myelinating oligodendrocytes among the endogenous cells.

To explore the human cell capacity of colonization of human the shiverer/Rag2^-/-^mouse spinal cord, cells were identified by immunolabeling of serial cross-sections with the human specific nuclear marker STEM101. Analysis at 8 wpg demonstrated the presence of exogenous cells not only within the dorsal funiculus adjacent to the injection site, but also in distant regions of both white and gray matter (Fig. 1). These findings indicate that grafted hiOLs not only successfully integrated the injection site but also contributed to extensive colonization of the neuraxis, encompassing both the ventral and dorsal spinal cord white and gray matter. Quantification revealed that STEM101⁺ cells extended over 4.81 ± 1.81 mm (n=5) at 8 wpg and 7.32 ± 2.19 mm (n=4) at 17 wpg (Fig. 1).

**Figure 1.**
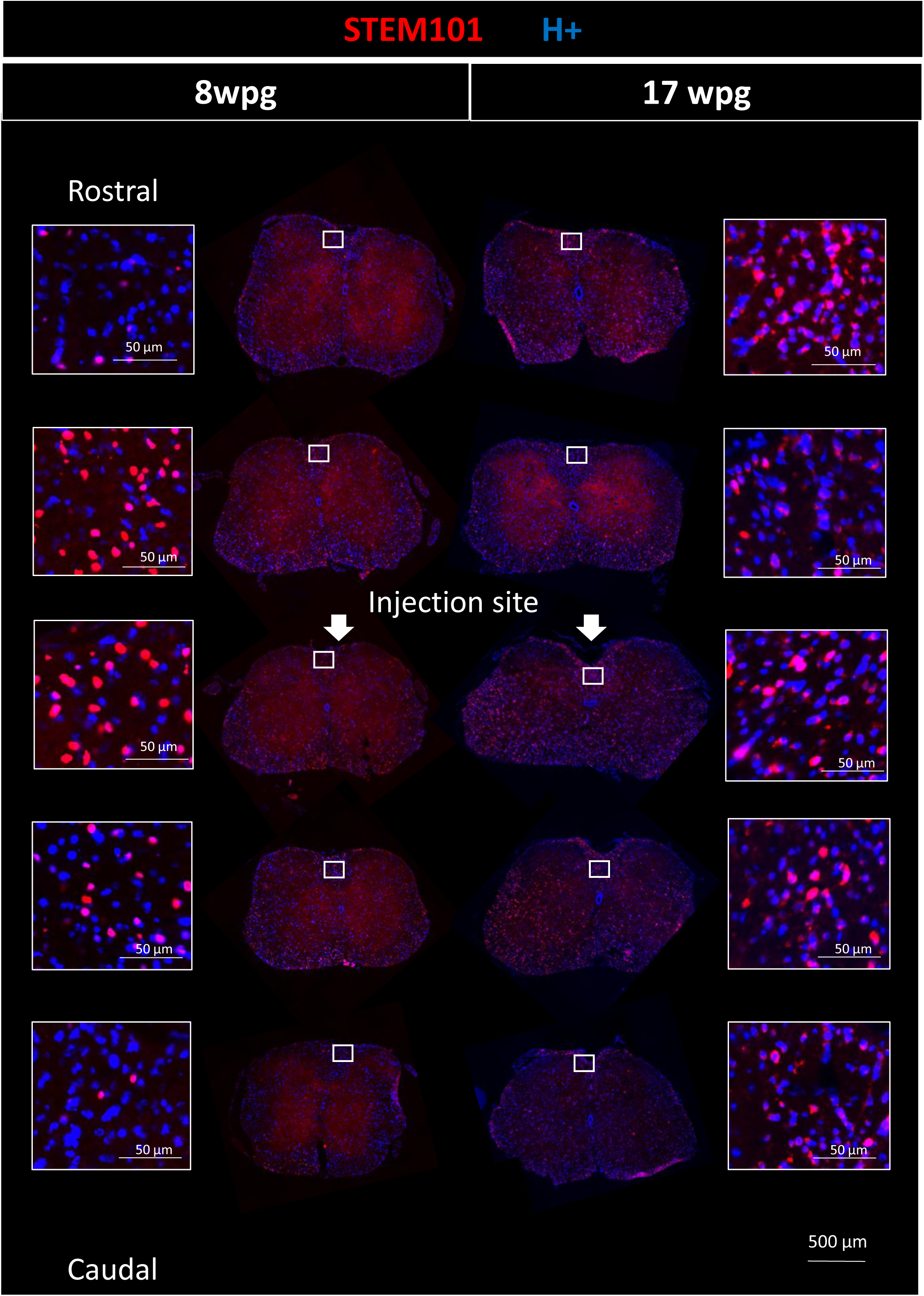
Serial coronal sections from the adult spinal cord of grafted Shi/Shi Rag2^−/−^ mice grafted at 3 weeks old and stained for the nuclear human marker STEM101 (red), show hiOLs presence throughout the entire neuraxis. Grafted hiOLs not only colonized the dorsal funiculus of the spinal cord, where the injection was performed, but also both ventral and dorsal white and gray matter. These images demonstrate the broad and timely rostro-caudal spread of human cells, spanning approximately 4.81 ± 1.81 mm (n = 5 mice) and 7.32 ± 2.19 mm (n=4) at 8 and 17 weeks post-grafting respectively. Scale bars, 500 and 50 μm.

To analyze the differentiation of the O4^+^ hiOL progenitors into mature oligodendrocyte, we quantified the proportion of human cells co-expressing the pan oligodendroglial marker OLIG2, the oligodendrocyte differentiation marker , CC1, together with STEM101 in the dorsal funiculus (Fig.2). At 8 wpg, nearly 40% of the grafted cells had already differentiated into mature oligodendrocytes (Fig. 2 K). To evaluate the temporal dynamic of differentiation, we further analyzed the exogenous cells at two later time points: 12 and 17 wpg (Fig.2 K). The proportion of mature CC1+OLIG2+STEM+ oligodendrocytes over total OLIG2+STEM+ cell population (Fig. 2 A, E and H, arrows) increased significantly by 12 wpg, with no further change at 17 wpg compared to 8 wpg (8wpg: 40.64 ± 2.75%, 12 wpg: 77.30 ± 1.01%, 17 wpg: 80.95 ± 1.88%) (8 wpg vs both 12 wpg and 17 wpg p<0.0001) (Fig. 2K). A group of grafted hiOL remained undifferentiated as adult-like hOPCs (CC1-OLIG2+STEM+) at all times (Fig. 2 A, E and H, arrow heads).

**Figure 2.**
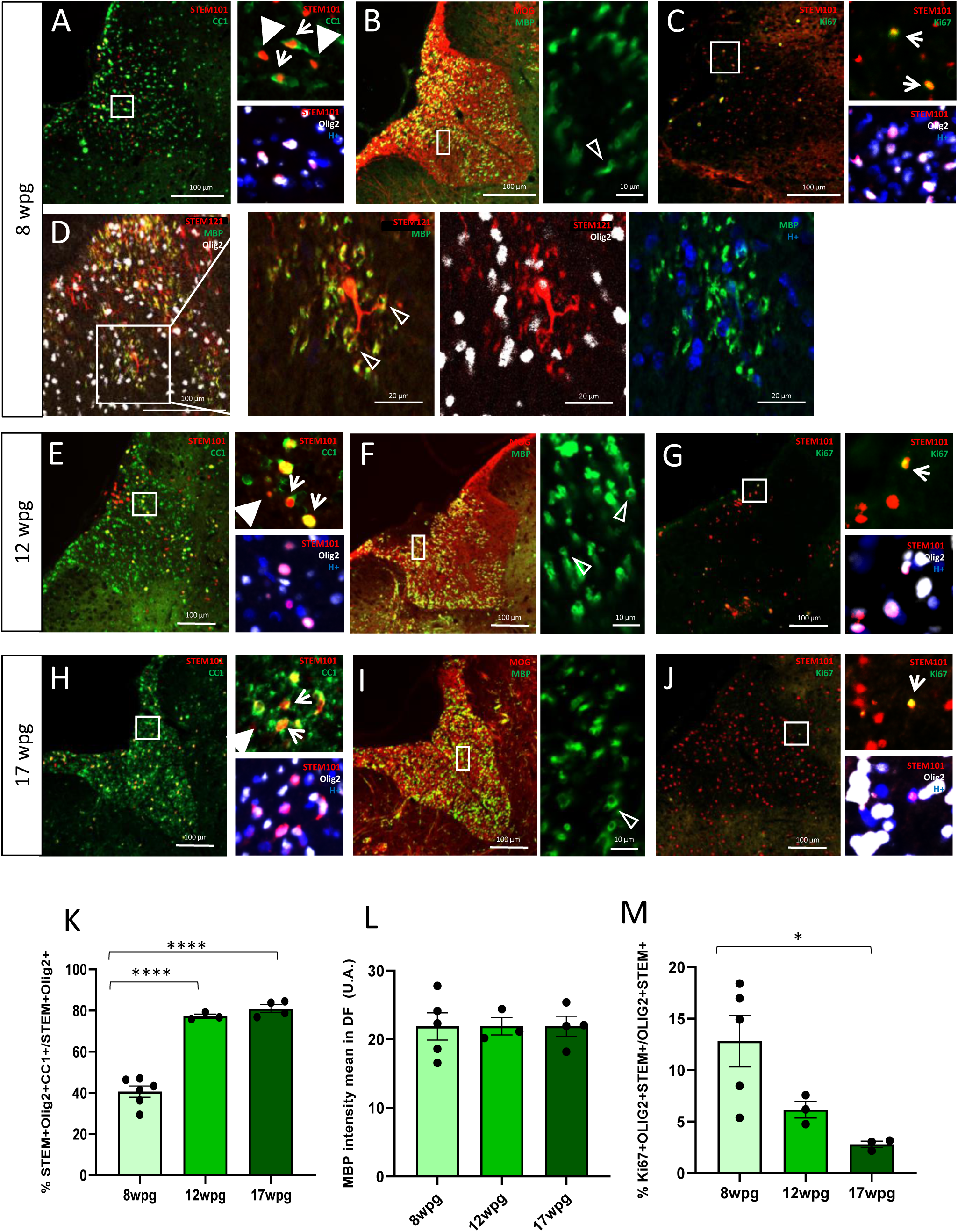
Engraftment of O4+ hiOL progenitors in the developing spinal cord leads (A-J). **A, E** and **H:** combined immunodetection of human nuclei marker STEM101 (red) with CC1 (green) and OLIG2 (white). Arrows depicting CC1+OLIG2+STEM+ while white arrow-heads point CC1-OLIG2+STEM+ and empty arrow-heads point MBP+ tubular/donut-like myelin structures. **B, F** and **I**: double immunostaining of host myelin (MOG, red) and grafted-derived myelin-like structures (MBP, green). **C, G** and **J:** combined immunodetection of human nuclei marker STEM101 (red) with Ki67 (green) and OLIG2 (white). **D**: combined immunostaining of human cytoplasmatic marker STEM121 (red), grafted-derived myelin-like structures (MBP, green) and OLIG2 (white). Analysis were performed at 8 (**A, B, C**), 12 (**E, F, G**), and 17 (**H, I, J**) wpg of CC1+ OLIG2+ STEM+ over OLIG2+ STEM+ cells (**K**), mean MBP intensity in the dorsal funiculus (A.U.) (**L**) and Ki67+ OLIG2+ STEM+ over OLIG2+ STEM+ cells (**M**). The percentage of mature OLs was significantly time regulated reaching a plateau at 12 wpg contributing to myelin formation while the percentage of human Ki67 cells was timely downregulated. One-way ANOVA followed by Tukey’s multiple comparison tests (*n* = 3 to 4 mice per group). Error bars represent SEMs. \**P* < 0.05 and \*\*\*\**P* < 0.0001. Scale bars, 100, 20 and 10 μm.

We further questioned whether the grafted O4+ hiOL cells differentiated into myelin-forming oligodendrocytes. To this aim, complementary immunostaining for MBP in conjunction with MOG was assessed. Since the host shiverer mice lack endogenous MBP, the presence of MBP indicates that it originates exclusively from grafted human cells, while MOG detects rodent and human myelin. MBP staining revealed MBP-positivity with some ring-shaped myelin-like structures within the colonized areas of the spinal cord at all time-points in the dorsal (Fig. 2 B, F, I and Fig. S1), and ventral white matter and even sometimes in the grey matter (Fig. S1). Combining detection of human cytoplasmic STEM 121 with MBP expressions revealed connection of these MBP+ rings to STEM+ processes reinforcing the human origin of the myelin-like structures (Fig.2D). Surprisingly, despite the significant increase in human oligodendrocyte maturation (Fig. 2K), the presence of donor-myelin did not increase with time in the dorsal funiculus (Fig. 2L) (mean MBP intensity: 8wpg: 21.88 ± 1.99, 12wpg: 21.92 ± 1.28, 17wpg: 21.91 ± 1.45 arbitrary units (A.U.)). MBP-positive myelin-like structures were observed along the same distance as the STEM+ nuclei (mean MBP distance:7.32 ± 2.19 mm (n=4)) at 17 wpg (Fig. S1), indicating that migration and myelination were tightly correlated.

Since a pool of exogenous human cells did not fulfill differentiation (CC1-OLIG2+STEM+, Fig. 2 A, E, H, arrow heads), the proliferative status of the grafted O4+ hiOL in engrafted spinal cords was analyzed by immunolabeling of Ki67, a nuclear marker of actively dividing cells. We found that a subset of exogenous oligodendroglial cells, expressing the human nuclear marker STEM101 in combination with OLIG2, expressed Ki67. This pool of proliferative human oligodendroglial cells indicates the presence of a reservoir of adult-like hOPCs that could respond to CNS needs. Although a low proportion of Ki67+OLIG2+STEM+ were observed at all times (3 to 13% of Ki67+OLIG2+STEM+ over STEM+OLIG2+ cells) (Fig. 2 C, G and J), a significant decrease in the proliferating exogenous population was noted at 17 wpg compared with 8 wpg (8 wpg: 12.82 ± 2.52%, 12 wpg: 6.17 ± 0.81%, 17 wpg: 2.78 ± 0.31%, 8 wpg vs 17 wpg p=0.022) (Fig. 2M) consistent with their increased differentiation.

### Myelin depletion in response to demyelination is followed by settled hiOL recruitment, activation and differentiation to restore myelin

The above findings indicate that the grafted hiOLs do not only contribute to long-term myelination but also maintain a population of proliferative progenitors, that could contribute to homeostasis and myelin repair. Given that both adult-like OPCs and surviving mature oligodendrocytes can participate in myelin repair (Duncan et al., 2018; Mezydlo et al., 2023), we enquired whether these two exogenous populations already settled in the spinal cord could respond to demyelination.

To this aim, focal demyelination was induced by LPC injection into the dorsal funiculus of the chimeric human-mouse spinal cord at 8 wpg (Fig. 3A-C). The 8 wpg time-point was chosen as the optimal time-point for the robust differentiation of the grafted O4^+^ hiOL progenitors into mature oligodendrocytes, the presence of donor myelin and the proliferation dynamic of the adult-like hOPC pool. Characterization of the lesion 1 week post lesion (wpl) (8 wpg+1 wpl) showed the focal loss of myelin, as evidenced by a significant decrease in host- and donor-MOG-positive myelin and donor-MBP+ myelin (Fig. 3 D) (mean MBP intensity: 8 wpg: 21.88 ± 1.99, 8 wpg + 1 wpl: 15.54 ± 1.60, arbitrary units (A.U.), p=0.048) (Fig. 3L). The reduction in MBP expression was associated with a significant reduction in mature oligodendrocytes within the lesion when comparing 8 wpg (40.11 ± 3.31%) with 8 wpg + 1 wpl (24.51 ± 5.63%) (p=0.039, Fig. 3K). The percentage of proliferating Ki67+ human oligodendroglial cells was only slightly reduced comparing 8 wpg with 8 wpg + 1 wpl (8 wpg: 12.82 ± 2.52%, 8 wpg + 1 wpl: 8.81 ± 0.97%, ns) (Fig. 3 F, M).

**Figure 3.**
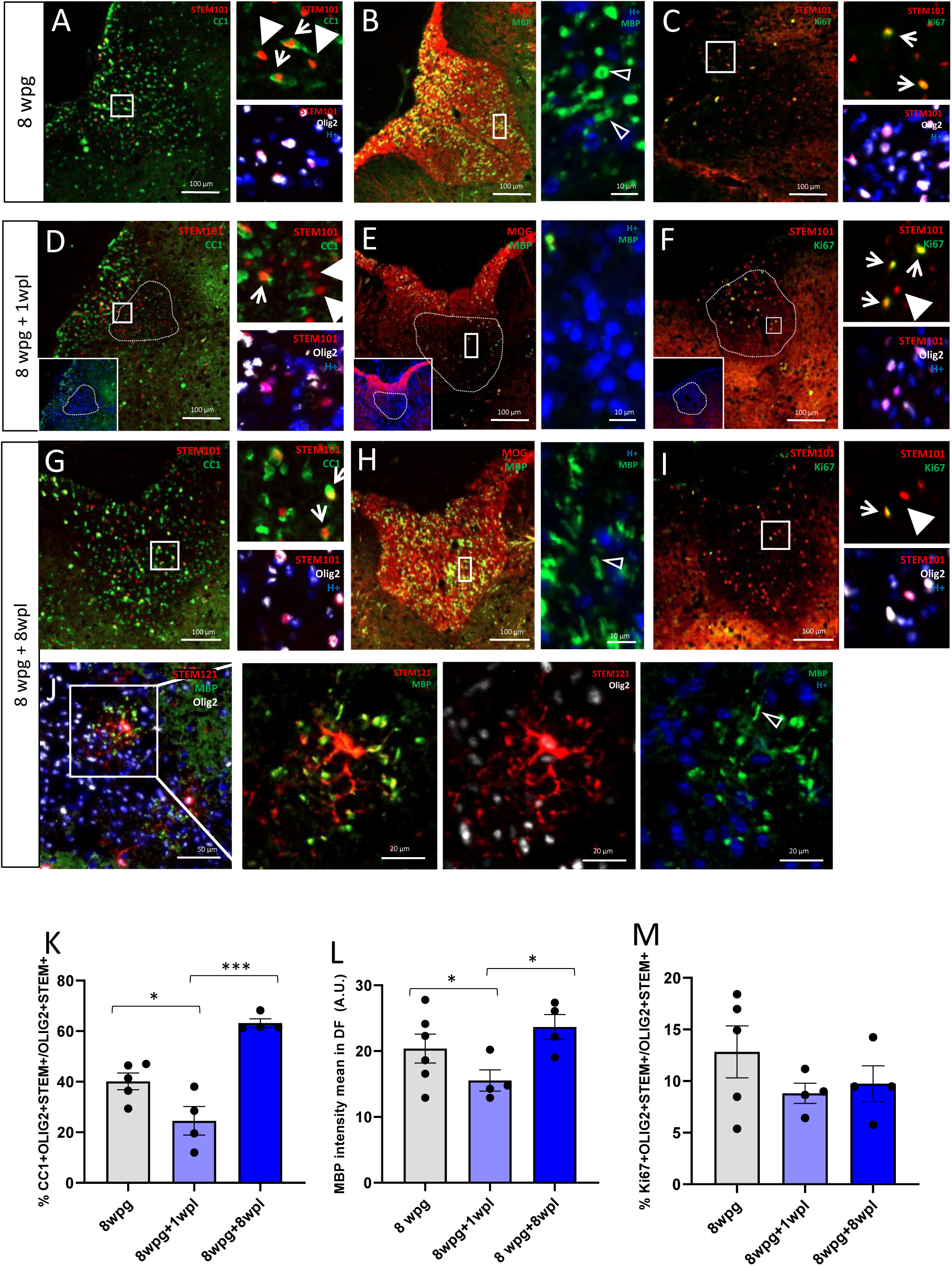
Regulation of hiOL numbers and their myelin in response to demyelination. **A, D** and **G:** combined immunodetection of human nuclei marker STEM101 (red) with CC1 (green) and OLIG2 (white). Arrows depicting CC1+OLIG2+STEM+ while arrow-heads point CC1-OLIG2+STEM+ and empty arrow-heads point MBP+ tubular/donut-like myelin structures. **B, E** and **H**: double immunostaining of host myelin (MOG, red) and grafted-derive myelin (MBP, green). **C, F** and **I:** combined immunodetection of human nuclei marker STEM101 (red) with Ki67 (green) and OLIG2 (white). **J**: combined immunostaining of human cytoplasmatic marker STEM121 (red), grafted-derive myelin (MBP, green) and OLIG2 (white). Analysis were performed at 8 (**A, B, C**), 12 (**D, E, F**), and 17 (**G, H, I**) wpg of CC1+ OLIG2+ STEM+ over OLIG2+ STEM+ cells (**K**), mean MBP intensity in the dorsal funiculus (A.U.) (**L**) and Ki67+ OLIG2+ STEM+ over OLIG2+ STEM+ cells (**M**). One-way ANOVA followed by Tukey’s multiple comparison tests (*n* = 4 to 6 mice per group). Error bars represent SEMs. \**P* < 0.05 and \*\*\**P* < 0.001. Scale bars, 100, 20 and 10 μm.

At 8 wpg + 8 wpl, expression of the donor MBP+ myelin recovered the initial values along with host/donor MOG+ myelin (Fig.3H, L) (mean MBP intensity: 8 wpg: 21.88 ± 4.44, vs 8 wpg + 8 wpl: 23.66 ± 1.89 A.U., *p*=0.017). Myelin restoration was associated with recovery of the donor mature oligodendrocyte population (Fig. 3K), with a significant 2.5-fold increase comparing 8 wpg + 1 wpl (24.51 ± 5.63%), with 8 wpg + 8 wpl (63.18 ± 1.68%) (p=0.0006, Fig. 3K), and a trend towards increase over the pre-lesion situation (8 wpg: 40.11 ± 3.31%). Analysis of STEM121+OLIG2+ population in the lesion, showed that some of the settled exogenous cells were associated with MBP+ myelin-like rings, pointing to their involvement in myelin restoration (Fig.3J). Results also suggested a consumption of the proportion of CC1-OLIG2+STEM+, likely adult-like OPCs, when comparing 1 wpl with 8 wpg. The proportion of proliferating donor oligodendroglial cells did not show any significant variation, just a trend towards reduction compared to 8 wpg and a minor increase compared to 1 wpl in agreement with the not significant reduction in immature hOL (CC1-OLIG2+STEM+, Fig. 2K, M and 3K, M).

These data strongly suggest participation of the settled exogenous mature oligodendrocytes to the process of myelin repair together with the maintenance of the exogenous human adult OPC-like reservoir.

### Testing the capacity to enhance the (re)myelination potential of human cells after myelin depletion

After demonstrating the capacity of the settled exogenous oligodendrocytes to be re-activated and to participate to the myelin repair process, we further questioned the susceptibility of the grafted hiOLs progeny to be boosted for remyelination by a pro-myelinating drug.

Focal demyelination was induced as above, in the humanized dorsal funiculus (Fig. 4). Bavisant, an orally active antagonist of the histamine H3 receptor, recently proven to enhance the pro-regenerative capacity of rodent and human OPCs (Gacem et al., 2026) was selected as potential candidate to enhance remyelination. Mice were orally treated with either vehicle (Fig. 4A-C) or bavisant (Fig. 4D-F) for eight weeks.

**Figure 4.**
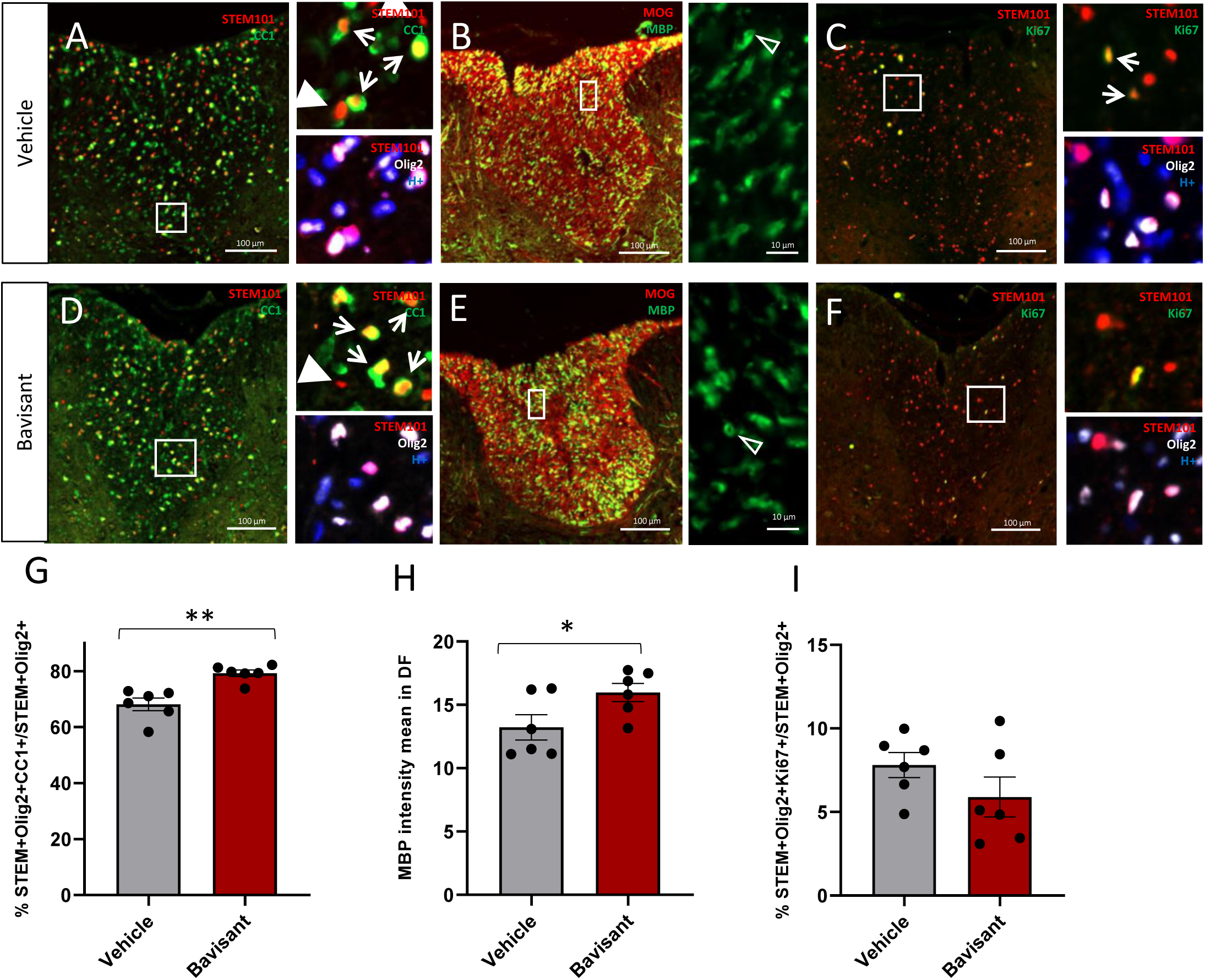
Differentiation of hiOL into remyelinating cells can be boosted by a pro-myelinating agent treatment. **A** and **D:** combined immunodetection of human nuclei marker STEM101 (red) with CC1 (green) and OLIG2 (white). Arrows depicting CC1+OLIG2+STEM+ while arrow-heads point CC1-OLIG2+STEM+ and empty arrow-heads point MBP+ tubular/donut-like myelin structures. **B** and **E**: double immunostaining of host myelin (MOG, red) and grafted-derived myelin (MBP, green). **C** and **F:** combined immunodetection of human nuclei marker STEM101 (red) with Ki67 (green) and OLIG2 (white). Analysis were performed at 8 wpg + 8wpl treated with vehicle (**A, B, C**), and bavisant (**D, E, F**) of CC1+ OLIG2+ STEM+ over OLIG2+ STEM+ cells (**G**), mean MBP intensity in the dorsal funiculus (A.U.) (**H**) and Ki67+ OLIG2+ STEM+ over OLIG2+ STEM+ cells (**I**). Unpaired t test (*n* = 6 mice per group). Error bars represent SEMs. \**P* < 0.05 and \*\**P* < 0.01. Scale bar, 100 μm.

Immunohistochemical analysis at 8wpl revealed substantial participation of the exogenous human oligodendrocytes in the repair process with the presence of CC1+OLIG2+STEM+ (Fig. 4A, D, arrows) and donor-derived MBP⁺ myelin-like structures (Fig. 4B, E) coexisting with host/donor-derived MOG⁺ myelin. While the vehicle group showed similar percentages of CC1+OLIG2+STEM+ over OLIG2+STEM+ cells to lesioned-untreated-mice at 8 wpg + 8 wpl (vehicle-treated mice: 68.12 ± 2.23%; untreated mice: 63.18 ± 1.68%) (Fig. 4G vs Fig. 3K), the proportion of CC1+OLIG2+STEM+ over OLIG2+STEM+ at 8 wpg+8 wpl was significantly increased in the bavisant-treated cohort (79.24 ± 1.33 %) compared to the vehicle cohort (68.12 ± 2.23%, p=0.001) (Fig.4G).

The same promoting effect was confirmed for donor MBP expression when comparing vehicle and bavisant treated groups (mean MBP+ intensity in vehicle-treated mice: 13.23 ± 0.99, bavisant-treated mice: 15.98 ± 0.71, arbitrary units (A.U.), p=0.049) (Fig. 4H). Although the MBP signal was lower than in the lesioned-untreated mice at 8 wpg + 8 wpl (mean MBP intensity: 23.66 ± 1.89 A.U.), it does not compromise the positive effect of the promyelinating agent bavisant.

Importantly, no significant difference was observed in the proliferation rates of the settled exogenous oligodendroglia between treated and non-treated mice (Fig. 4I) (vehicle-treated mice: 7.80 ± 0.74%, bavisant-treated mice: 5.89 ± 1.19%) indicating that bavisant primarily enhanced hiOL capacity to differentiate in myelin-like forming cells rather than promoting hiOL derived adult-like OPC proliferation.

### Bavisant enhances hiOLs remyelination of previously denuded host axons

Axons remyelinated by shiverer oligodendrocytes have incomplete and uncompacted myelin sheaths, while those remyelinated by wild-type murine or human oligodendrocytes have compact myelin. Ultrastructural analysis of lesion areas revealed the presence of unmyelinated axons, as well as remyelinated axons by murine (rare and uncompacted) and human (thin and compacted) oligodendrocytes in both vehicle and bavisant conditions. In Figure 5, representative electron micrographs illustrate these findings. The quantification of remyelination revealed a significant increase in remyelinated axons in bavisant-treated mice compared to the control group (Fig. 5G, H). This effect was significant when comparing the percentage of remyelinated axons to total axons (denuded or remyelinated) (Fig. 5G) (vehicle-treated mice: 7.097 ± 0.631, bavisant-treated mice: 14.591 ± 2.091, p=0.031) or to total number of remyelinated axons (Fig. 5H) (vehicle-treated mice: 22.07 ± 1.63, bavisant-treated mice: 47.61 ± 7.97, p=0.04). These results indicate that bavisant enhances remyelination by human oligodendrocytes and provides strong support for its potential as a therapeutic candidate for promoting CNS remyelination in humans.

**Figure 5.**
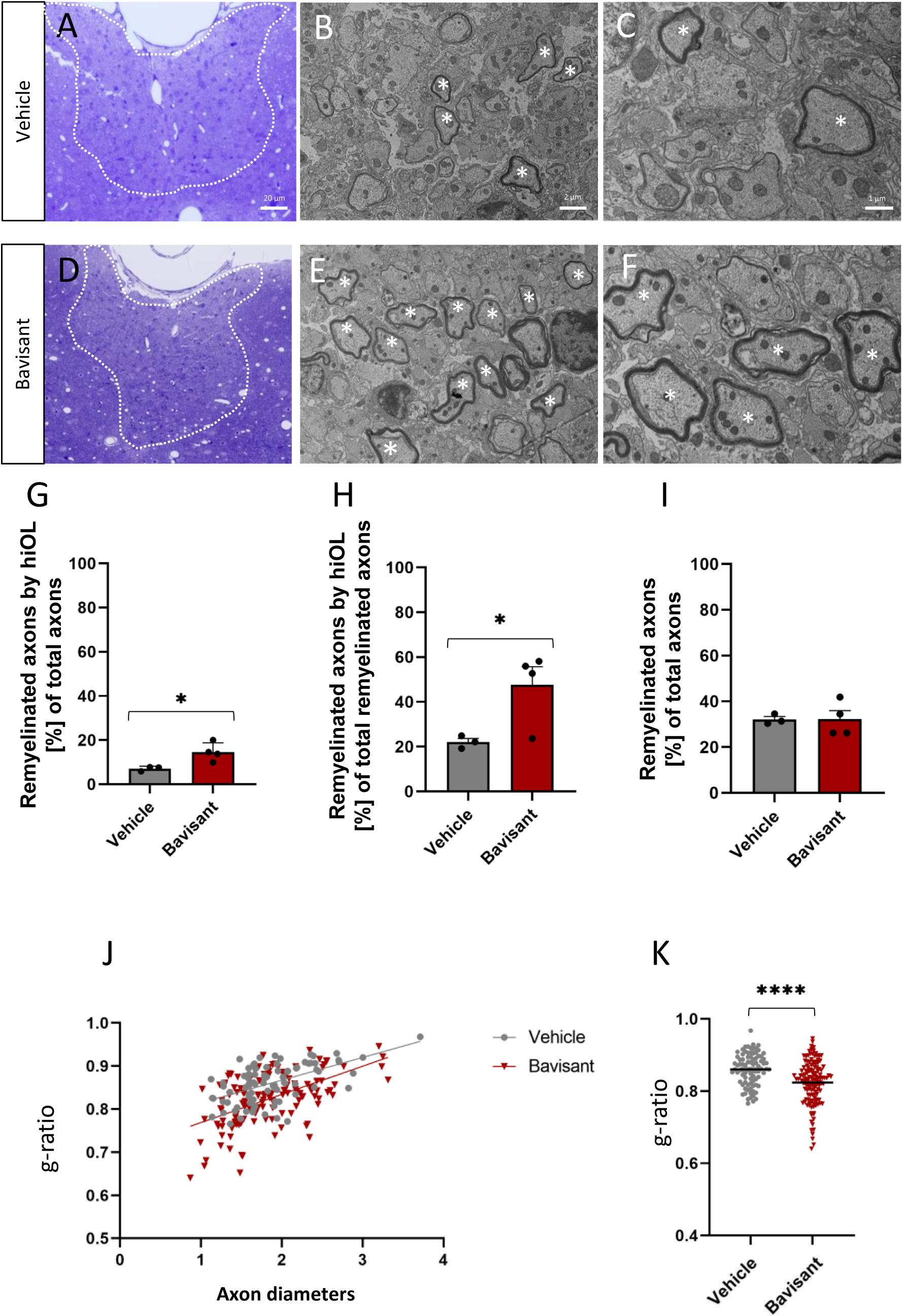
Ultrastructural validation of a novel remyelination paradigm by human hiOL and of bavisant efficacy to boost this process. **A, D**: Representative semi-thin sections of the dorsal funiculus (dashed line) stained with toluidine blue depicting hiOLs within LPC lesions in vehicle- (**A**) or bavisant- (**D**) treated groups. **B,C,E,F**: electron micrographs illustrating the presence of compact myelin (*) in spinal cord lesions of shiverer mice after grafted of hiOLs In vehicle treated animals, sparse myelinated axons are observed (**B** and **C**) In contrast, animals receiving bavisant exhibit a higher density of compact myelin (**E** and **F**). **G-I**: the percentage of axons remyelinated by hiOLs was calculated relative to the total number of axons within the analyzed area (**G**) and to the total remyelinated axons (**H**). quantification of the total number of remyelinated axons derived from either human cells or endogenous mouse oligodendrocytes was performed at 8 weeks post lesion following LPC induced demyelination in shiverer mice (**I**). **J-K**: Distribution of g-ratio (**J**) and g-ratio measurements (**K**) reveal a significant reduction in values following bavisant treatment, reflecting increased myelin sheath thickness. Scale bars, 20, 2 and 1 μm. Error bars represent SEMs. Significance was analyzed by unpaired t test *p< 0,05,****p< 0,0001.

Interestingly, the total number of remyelinated axons originating from combined exogenous human cells and endogenous murine oligodendrocytes did not differ significantly between groups (Fig. 5I) (vehicle-treated mice: 32.10 ± 1.25, bavisant-treated mice: 32.17 ± 3.80). This suggests that approximately 30% of axons undergo remyelination in shiverer mice at this stage in both conditions, and that human cells compete over endogenous cells to further enhance axonal remyelination under bavisant treatment. Moreover, analysis of the g-ratio reflecting myelin thickness revealed a significant increase of compact myelin sheath thickness shown as a decrease in g-ratio, in mice treated with bavisant compared to control (Fig. 5J, K) (p<0.0001) thus inmplaying that bavisant treatment increased the number of human myelin wraps around host axons.

## Discussion

Intrinsic mechanisms governing human oligodendrocytes differentiation display species-specific properties, which cannot be fully recapitulated in existing animal models. Moreover, in vitro and ex vivo approaches offer partial insights, while lacking the complex 3D in vivo environment that influences myelination. Therefore, there is a critical need for humanized in vivo models to accurately study human oligodendrocyte development and the impact of genetic and epigenetic alterations and/or external environmental cues on their capacity of myelination and repair.

Although previous studies have attempted to address these gaps (Ehrlich et al., 2017; Mozafari et al., 2020; Windrem et al., 2020), the translational relevance of these approaches remains limited. In this context, the use of human iPSC-derived cells (Ehrlich et al., 2017; Mozafari et al., 2020), rather than those from fetal origin (Levy et al., 2022; Windrem et al., 2020) to humanize animal models, offers the advantage of integrating patient-specific genetic information within an in vivo environment. This aspect is particularly relevant, as iPSC-progenitors can be generated directly from patients with diverse demyelinating or dysmyelinating disorders, thereby preserving their individual genetic backgrounds. Such an approach provides a unique opportunity to model patient-specific disease mechanisms and to evaluate promyelinating therapeutic responses in vivo.

We have demonstrated previously the ability of exogenous hiPSC-derived OPCs to colonize and myelinate the brain and spinal cord,and remyelinate the spinal cord following induced demyelination (Ehrlich et al., 2017; Mozafari et al., 2020). Yet, the capacity of the iPSC oligodendroglia to be challenged following their initial developmental myelinationhas not been established. Importantly, while in the demyelination-then-grafting paradigm, the myelin formed by the human cells is referred to as “remyelination” from the axon perspective (since axons have already been myelinated prior subsequent demyelinated), the grafted human cells should be considered as embryonic cells (as opposed to settled adult cells) since, they never produced myelin before and therefore, represent the first cycle of de novo myelination.

An additional critical consideration is the establishment of a translationally relevant humanized animal model that ensures the donor origin of the newly formed myelin over the host one. For this reason, the use of MBP-deficient shiverer mice is indispensable. Furthermore, although both myelination and remyelination involve the generation of myelin, the underlying mechanisms differ substantially (Moyon et al., 2017). Developmental myelination and remyelination occur within distinct cellular and molecular contexts, particularly in diseased or inflamed environments (Hill et al., 2025), where remyelination must take place along previously demyelinated axons or be carried out by oligodendroglial cells that have undergone aging prior demyelinating insults. Nevertheless, despite the necessity of immunodeficient mice for humanized animal models, the influence of lymphocytes on oligodendrocyte maturation and regeneration in the Rag2^-/-^ strain should not be underestimated. Taking into account these various considerations, we have developed and characterized a humanized mouse model by transplanting hiOLs into the developing spinal cord of MBP-deficient and immunodeficient shiverer/Rag2^⁻/⁻^ mice. Longitudinal analysis of the developing spinal cord, demonstrated that after one single spinal cord injection, the grafted cells migrated efficiently and timely colonized the host spinal parenchyma, invading both the dorsal and ventral white and gray matter regions. Moreover, the human iPSC-derived progeny exhibited timely and progressive differentiation into mature oligodendrocytes with MBP+ myelin-like features. In addition to the generation of mature oligodendrocytes, a proliferating pool of human adult OPCs was present at all time points studied. While the number of these cells decreased overtime, their sustained presence supports the existence of a stable human progenitor reservoir that could be available, in addition to mature oligodendrocytes (exogenous and endogenous), for subsequent myelin repair, as observed in other human-mouse chimeric mouse models (Ehrlich et al., 2017; Levy et al., 2022; Windrem et al., 2020). These developmental observations recapitulate important features of CNS biology, including the asynchronous and heterogeneous maturation of OPCs available to respond to CNS plasticity, demyelinating insults or promyelinating agents.

The extensive colonization by exogenous human cells of the host spinal cord provided the opportunity to model both developmental myelination and remyelination within the same animal model and with the same grafted human cell population. To this end, we induced focal demyelination in the chimeric adult dorsal spinal cord to challenge the fate and response of the settled exogenous human cells in response to this insult. This model offers several advantages over the classical approaches, such as cell engraftment into demyelinating lesions or models limited to humanized developmental myelination alone.

First, the use of genetically hypomyelinated mice enabled to demonstrate the capacity of human oligodendrocytes to function as remyelinating cells in vivo, and distinguishable from host myelin by their MBP⁺ expression and the presence of multilayered axonal wrapping as seen by electron microscopy.

Second, it opens the opportunity to mimic features of endogenous remyelination in which iPSC-derived oligodendrocytes and OPCs that have settled in the adult spinal cord, are challenged by demyelination as opposed to embryonic-derived populations. Exogenous oligodendroglial cells responded to LPC-induced focal demyelination with a transient reduction in the number of mature OLs and MBP^+^ myelin, followed by the re-establishment of the oligodendroglial population and restoration of donor-derived myelin over time. Interestingly, the proportion of mature oligodendrocytes recovered after demyelination to levels achieved prior to the lesion, although it remained lower than in non-injured conditions. Consistently, MBP expression after demyelination recovered to levels achieved both before lesion (at 8 wpg) and in naïve mice (at 17 wpg), suggesting a maximal myelination capacity achievable in the shiverer spinal cord.

Third, analysis of the proliferative activity of the exogenous cells, indicated the persistence of an adult OPC-like reservoir that can be activated in response to demyelination and can potentially contribute with the endogenous and exogenous mature oligodendrocytes to CNS plasticity and repair. Further studies involving repeated demyelinating insults could help to clarify the dynamics and resilience of this hiOL-OPC reservoir. Moreover, clarifying whether it is one or both of these populations that can participate in repair under pathological conditions, would constitute the scope of future translationally relevant investigation to specifically target one of these populations.

In addition, the present study further demonstrates the translational potential of this model in evaluating the therapeutic efficacy of bavisant, a histamine H3 receptor antagonist previously reported to promote remyelination in rodent models of demyelination (Gacem et al., 2026). Bavisant treatment of the demyelinated humanized spinal cords led to a significant increase in the proportion of human-derived mature oligodendrocytes and MBP^+^ myelin respect to vehicle, as well as an increased proportion of remyelinated axons, without altering the proliferation capacity of the exogenous cells. These data suggest that bavisant enhances the differentiation and myelination capacity of hiOLs rather than acting on OPC proliferation, highlighting its potential as a remyelinating therapeutic in human disease contexts. It is worth noting that although the promyelinating effect of bavisant is obvious in our paradigm, the overall MBP signal was significantly lower after bavisant or vehicle treatment compared with the untreated condition. Despite this unexpected outcome, the continuous gavage administration of the vehicle and compound over an eight-week period may have induced stress that compromised myelin formation. Although this hypothesis requires further investigation, adopting voluntary oral administration methods would represent an ethical and reasonable consideration for future drug-testing studies.

Electron microscopy analysis provided ultrastructural confirmation of the donor-derived remyelination in vehicle and bavisant groups and showed improved g-ratios in bavisant-treated animals, further validating the functional integration of hiOLs as remyelinating cells. Moreover, the present results not only confirm bavisant as an effective and rapid promyelinating compound for remyelinating demyelinated axons but also demonstrate the availability of a novel protocol to address human-derived remyelination. Compared to our prior study (Gacem et al., 2026), when hiOLs were grafted within the lesion, the re-myelination ratio in control groups increased from 10% (as reported in the first study with bavisant) to 20% in the current humanized model. Specifically, this experimental paradigm of introducing demyelination after maturation of graft-derived oligodendrocytes shows that hiOL-progeny achieves a higher baseline level of remyelination than when hiOLs were grafted after LPC lesion (Gacem et al., 2026). This might be due to the slow tempo of differentiation of the human cells grafted as “embryonic-like” OPCs compared to exogenous human cells already available as resident mature oligodendrocytes and adult-like OPCs as used in the present model. This enhancement in remyelination, facilitated by the presence of already settled human cells, improves the sensitivity of the assay, allowing for clearer differentiation between experimental groups and providing a more robust model for testing the effects of remyelinating molecules in human cells. This feature boosts the model’s sensitivity to detect pro-remyelinating effects in potential therapeutic agents and thus refining its utility for drug screening.

Overall, this novel paradigm offers a robust and translationally relevant platform for studying the mechanisms of hiOL maturation, myelination, and remyelination following CNS injury. These findings underscore the value of a humanized oligodendroglia chimeric mouse model to delineate the molecular and cellular processes governing myelination and remyelination in a human genetic context. The model holds substantial promise not only for studying disease-specific oligodendrocyte dysfunction but also for the preclinical evaluation of myelin repair therapies across a range of demyelinating and neurodegenerative disorders. Future work will benefit from incorporating patient-derived iPSCs to investigate pathological variants, as well as employing high-throughput and single-cell profiling approaches to unravel the mechanistic underpinnings of impaired myelination in disease.

## Acknowledgments

We thank the Paris Brain Institute Cell Culture (CELIS), Histology (Histomics), Imaging (ICMQuant), Genotyping and sequencing core facility (iGenSeq), and Rodent (PhenoParc) facilities for the technical advice. We thank N. Sarrazin and J. Droesbeke for the help with in vivo experiments. We are grateful to the Pitié-Salpêtrière Hospital FAC S facility for cell sorting.

## References

Adams, K.L., et al., 2021. Intrinsic and extrinsic regulators of oligodendrocyte progenitor proliferation and differentiation. Semin Cell Dev Biol. 116, 16–24.

Barba-Reyes, J.M., et al., 2025. Oligodendroglia vulnerability in the human dorsal striatum in Parkinson’s disease. Acta Neuropathol. 149, 46.

Baydyuk, M., et al., 2020. Extrinsic Factors Driving Oligodendrocyte Lineage Cell Progression in CNS Development and Injury. Neurochem Res. 45, 630–642.

Blanchard, B., et al., 2013. Tocopherol derivative TFA-12 promotes myelin repair in experimental models of multiple sclerosis. J Neurosci. 33, 11633–42.

Boulanger, J.J., Messier, C., 2014. From precursors to myelinating oligodendrocytes: contribution of intrinsic and extrinsic factors to white matter plasticity in the adult brain. Neuroscience. 269, 343–66.

Buchet, D., et al., 2011. Human neural progenitors from different foetal forebrain regions remyelinate the adult mouse spinal cord. Brain. 134, 1168–83.

Chanoumidou, K., et al., 2020. Stem cell derived oligodendrocytes to study myelin diseases. Glia. 68, 705–720.

Chen, X., et al., 2018. Genetic and Epigenetic Alterations Underlie Oligodendroglia Susceptibility and White Matter Etiology in Psychiatric Disorders. Front Genet. 9, 565.

Depp, C., et al., 2023. Myelin dysfunction drives amyloid-beta deposition in models of Alzheimer’s disease. Nature. 618, 349–357.

Dimovasili, C., et al., 2023. Aging compromises oligodendrocyte precursor cell maturation and efficient remyelination in the monkey brain. Geroscience. 45, 249–264.

Duncan, I.D., et al., 2018. The adult oligodendrocyte can participate in remyelination. Proc Natl Acad Sci U S A. 115, E11807–E11816.

Ehrlich, M., et al., 2015. Distinct Neurodegenerative Changes in an Induced Pluripotent Stem Cell Model of Frontotemporal Dementia Linked to Mutant TAU Protein. Stem Cell Reports. 5, 83–96.

Ehrlich, M., et al., 2017. Rapid and efficient generation of oligodendrocytes from human induced pluripotent stem cells using transcription factors. Proc Natl Acad Sci U S A. 114, E2243–E2252.

Falcao, A.M., et al., 2018. Disease-specific oligodendrocyte lineage cells arise in multiple sclerosis. Nat Med. 24, 1837–1844.

Falkai, P., et al., 2023. Disturbed Oligodendroglial Maturation Causes Cognitive Dysfunction in Schizophrenia: A New Hypothesis. Schizophr Bull. 49, 1614–1624.

Filley, C.M., 2021. Cognitive Dysfunction in White Matter Disorders: New Perspectives in Treatment and Recovery. J Neuropsychiatry Clin Neurosci. 33, 349–355.

Fodder, K., et al., 2023. The contribution of DNA methylation to the (dys)function of oligodendroglia in neurodegeneration. Acta Neuropathol Commun. 11, 106.

Gacem, N., et al., 2026. In silico screening and preclinical validation identify bavisant as a therapeutic candidate for multiple sclerosis. Sci Transl Med. 18, eads0633.

Garcia-Leon, J.A., et al., 2020. Generation of oligodendrocytes and establishment of an all-human myelinating platform from human pluripotent stem cells. Nat Protoc. 15, 3716–3744.

Han, S., et al., 2022. Functions and dysfunctions of oligodendrocytes in neurodegenerative diseases. Front Cell Neurosci. 16, 1083159.

Hill, B.M., et al., 2025. Monocyte-secreted Wnt reduces the efficiency of central nervous system remyelination. PLoS Biol. 23, e3003073.

Horiuchi, M., et al., 2024. ALS-linked mutant TDP-43 in oligodendrocytes induces oligodendrocyte damage and exacerbates motor dysfunction in mice. Acta Neuropathol Commun. 12, 184.

Howng, S.Y., et al., 2010. ZFP191 is required by oligodendrocytes for CNS myelination. Genes Dev. 24, 301–11.

Hughes, E.G., Appel, B., 2016. The cell biology of CNS myelination. Curr Opin Neurobiol. 39, 93–100.

Jeffries, M.A., et al., 2021. mTOR Signaling Regulates Metabolic Function in Oligodendrocyte Precursor Cells and Promotes Efficient Brain Remyelination in the Cuprizone Model. J Neurosci. 41, 8321–8337.

Kang, S.H., et al., 2013. Degeneration and impaired regeneration of gray matter oligodendrocytes in amyotrophic lateral sclerosis. Nat Neurosci. 16, 571–9.

Levy, M.J.F., et al., 2022. High Dose Pharmaceutical Grade Biotin (MD1003) Accelerates Differentiation of Murine and Grafted Human Oligodendrocyte Progenitor Cells In Vivo. Int J Mol Sci. 23.

Lopez-Muguruza, E., Matute, C., 2023. Alterations of Oligodendrocyte and Myelin Energy Metabolism in Multiple Sclerosis. Int J Mol Sci. 24.

Luo, J.X.X., et al., 2022. Human Oligodendrocyte Myelination Potential; Relation to Age and Differentiation. Ann Neurol. 91, 178–191.

Mezydlo, A., et al., 2023. Remyelination by surviving oligodendrocytes is inefficient in the inflamed mammalian cortex. Neuron. 111, 1748–1759 e8.

Mot, A.I., Depp, C., Nave, K.A., 2018. An emerging role of dysfunctional axon-oligodendrocyte coupling in neurodegenerative diseases. Dialogues Clin Neurosci. 20, 283–292.

Moyon, S., et al., 2017. Efficient Remyelination Requires DNA Methylation. eNeuro. 4.

Mozafari, S., et al., 2020. Multiple sclerosis iPS-derived oligodendroglia conserve their properties to functionally interact with axons and glia in vivo. Sci Adv. 6.

Orthmann-Murphy, J., et al., 2020. Remyelination alters the pattern of myelin in the cerebral cortex. Elife. 9.

Patel, J.R., et al., 2010. CXCR4 promotes differentiation of oligodendrocyte progenitors and remyelination. Proc Natl Acad Sci U S A. 107, 11062–7.

Philips, T., Rothstein, J.D., 2017. Oligodendroglia: metabolic supporters of neurons. J Clin Invest. 127, 3271–3280.

Reinhardt, P., et al., 2013. Derivation and expansion using only small molecules of human neural progenitors for neurodegenerative disease modeling. PLoS One. 8, e59252.

Rey, F., et al., 2024. Role of epigenetics and alterations in RNA metabolism in leukodystrophies. Wiley Interdiscip Rev RNA. 15, e1854.

Schneider, C.A., Rasband, W.S., Eliceiri, K.W., 2012. NIH Image to ImageJ: 25 years of image analysis. Nat Methods. 9, 671–5.

Schuster, K.H., et al., 2022. Impaired Oligodendrocyte Maturation Is an Early Feature in SCA3 Disease Pathogenesis. J Neurosci. 42, 1604–1617.

Sim, F.J., et al., 2002. The age-related decrease in CNS remyelination efficiency is attributable to an impairment of both oligodendrocyte progenitor recruitment and differentiation. J Neurosci. 22, 2451–9.

Stadelmann, C., et al., 2019. Myelin in the Central Nervous System: Structure, Function, and Pathology. Physiol Rev. 99, 1381–1431.

Tsarouchas, T.M., et al., 2025. Protocol for assessing myelination by human iPSC-derived oligodendrocytes in Shiverer mouse ex vivo brain slice cultures. STAR Protoc. 6, 103609.

Windrem, M.S., et al., 2020. Human Glial Progenitor Cells Effectively Remyelinate the Demyelinated Adult Brain. Cell Rep. 31, 107658.

Xu, H., et al., 2020. m(6)A mRNA Methylation Is Essential for Oligodendrocyte Maturation and CNS Myelination. Neuron. 105, 293–309 e5.

