## Supplemental Figure 1 for "Development of a Humanized Mouse Model for Studying adult Spinal Cord myelination, remyelination and Drug Efficacy"

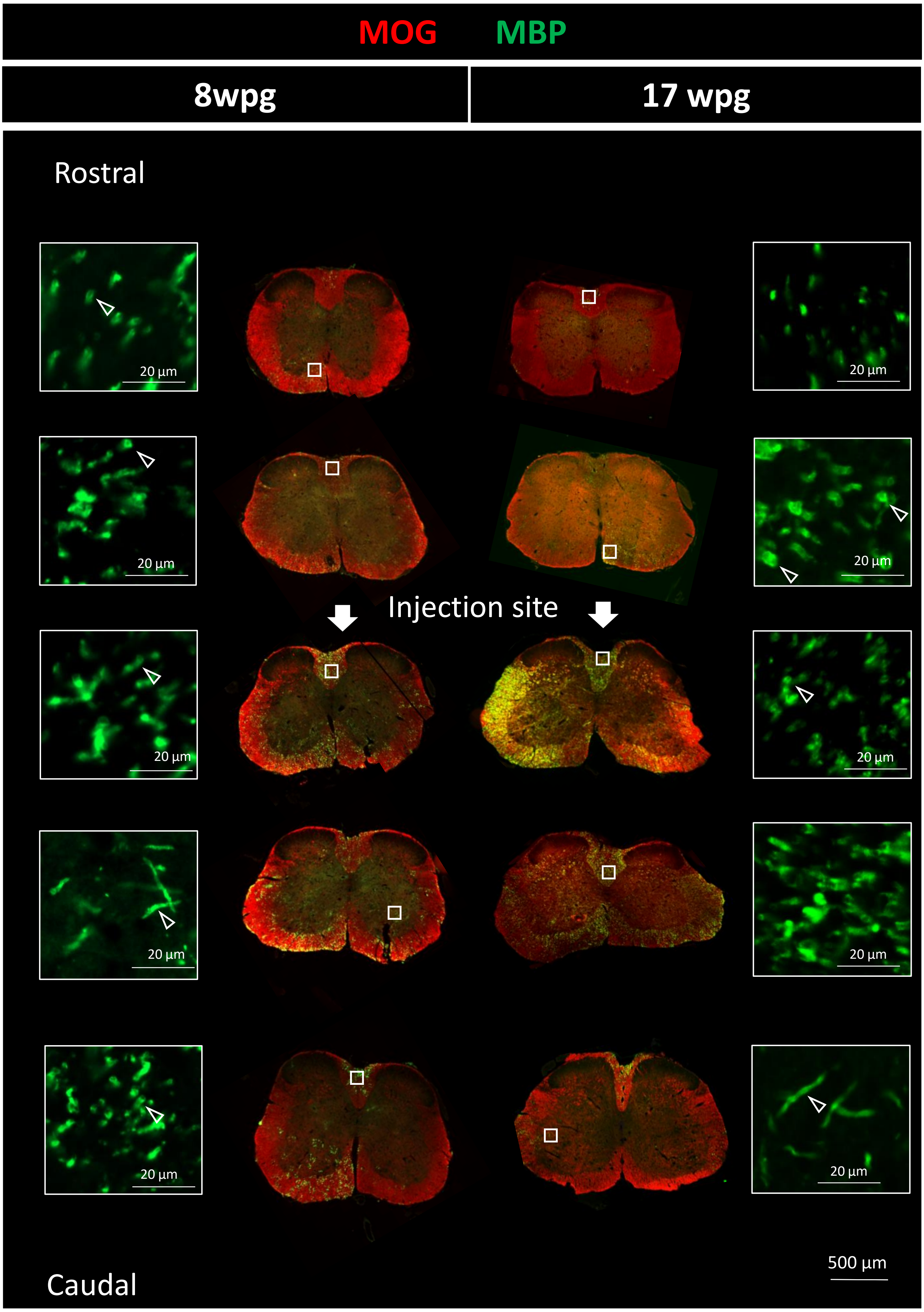


Figure S1. Human myelin is extensively distributed and integrates efficiently with endogenous myelin in adult mice when grafted at 3-weeks of age. Serial coronal sections from the spinal cord of grafted adult Shi/Shi Rag2$^{-/-}$ mice stained for MOG (red), and MBP (green) show hiOLs presence, MOG+/MBP− endogenous myelin and MOG+/MBP+ human-derived myelin. MBP+ with tubular or donut-like myelin structures (empty arrow-heads in insets). These images demonstrate the broad rostro-caudal spread of MBP+ human myelin, spanning approximately 4.81 ± 1.81 mm, at 8 wpg (n = 5 mice) and 7.32 ± 2.19 mm (n=4) at 17 wpg. Scale bars, 500 and 20 µm.
